# Nucleosome sensitivity distinguishes colon polyps based on their transformation status

**DOI:** 10.1101/2022.11.08.515478

**Authors:** Mahdi Khadem, Kimberlee Kossick, Yaroslav Fedyshyn, Lisa Boardman, Jonathan H. Dennis, Brooke R. Druliner

## Abstract

One of the keys to eliminating the personal and financial costs of cancer lies in the early detection of the disease. Consequently, effective cancer interventions increasingly rely on our understanding of the earliest cellular and nuclear events that lead to oncogenic transformation. Colorectal cancer, the third most prevalent cancer in the United States, results from the transformation of polyps. Our group demonstrated that the alteration of chromatin organization is a pivotal event in this oncogenic transformation. Here, we analyze the differences of the nucleosomal sensitivity to mocroccocal nuclease (MNase) between histopathologically matched pre-cancerous polyps taken from patients that did not develop cancer (cancer-free polyps, CFP) and those that did develop cancer (cancer-associated polyps, CAP). We produced high-resolution nucleosome distribution and nucleosome sensitivity maps from each of the five CFP patient samples and three CAP patient samples. We show that nucleosome distribution is largely invariant between CFP and CAP samples. Nucleosome sensitivity, however, is a powerful analysis that can identify genomic locations that distinguish CFP from CAP. We have identified more than 1000 genomic locations with altered nucleosomal sensitivity that discriminate between CAP and CFP. Furthermore, we show that these genomic locations with altered nucleosomal sensitivity between CFP and CAP include genes that play critical roles in oncogenic transformation. We propose that nucleosome sensitivity serves as a robust biomarker indicating the oncogenic potential of precancerous polyps and could be used for the early detection of polyps that will become cancerous.

## INTRODUCTION

Colorectal cancer (CRC) is the third most common cancer and the second leading cause of cancer death for men and women combined in the United States (1). CRC develops through a multifactorial process that begins with abnormal cellular growth of the colon epithelium, which can develop into a polyp and then potentially transform to cancer (2,3). The pathways involved in the transformation of a polyp to CRC are understood in the polyp to cancer sequence, but the mechanisms underlying why one polyp transforms while another doesn’t, are not understood (3–7). Understanding the mechanisms involved in what permits the transition state from benign growth (polyp) to cancer is central to developing interventions for targeted early detection or individualized clinical management.

Eukaryotic genomes consist of chromatin, which is DNA packaged together with histone proteins into nucleosomes. A hallmark of the transformed phenotype is altered chromosome structure, and important studies have implicated classes of chromatin regulatory complexes in cellular transformation. Inappropriate regulation of chromatin structure may inhibit normal cell development and differentiation and may represent the origin of cellular transformation. Cellular transformation is driven by the interplay of genetic and epigenetic changes, and a large body of research has identified modifications of chromatin and the role of those modifications in regulating genes. We have previously identified telomere, genomic, transcriptomic, and methylation changes that distinguish polyps that have transformed to cancer and those that have not (8). We examined these features in cancer-adjacent polyps (CAPs) and cancer-free polyps (CFPs), which are polyps that are clinically and histologically indistinguishable. In a CAP, the cancer which arose from the polyp is present while CFPs have no adjacent cancer.

Over the past several decades extensive research has been done to further understand the relationship between epigenetics and the development of cancer. These studies have determined factors that are implicated in cancer including DNA methylation, histone modifications, chromatin remodelers, and most recently, our work on nucleosome distribution alterations in early cancer (9,10). We have previously shown that nucleosome distribution is an early, widespread transformation event in lung and colon adenocarcinoma. In this current study, we map nucleosome distribution and sensitivity in polyps. Here, we identify chromatin features of polyps based on different clinical outcomes, and thus provide clinically relevant molecular subtypes of polyps based on chromatin structure. The location and density of nucleosomes with respect to the underlying DNA sequence is an important factor determining access to the genome for DNA-templated processes.

## RESULTS

### Nucleosome distribution is invariant between cancer-free and cancer-associated polyps

The distribution of nucleosomes along the genome likely plays a role in the accessibility of the genome to regulatory factors (11). Because we previously reported widespread nucleosome distribution changes as a feature of early stage colorectal cancers, we wanted to determine the signature of nucleosome organization around transcription start sites in precursor lesions to cancer, polyps. To that end, we studied eight polyp samples from patients with differing outcomes: polyps that were not associated with cancer, and other, clinically indistinguishable polyps that were associated with cancer at the time of detection. These polyps were categorized into two groups: cancer-adjacent polyps (CAPs), derived from patients with colon adenocarcinoma and cancer free polyps (CFPs), derived from patients with no observed malignant tumors. We used our MNase-Transcription Start Site Sequence Capture method (mTSS-seq) established previously to produce high-resolution maps of lung adenocarcinoma (9) and colorectal carcinoma (10). Our mTSS-seq combines in-solution targeted enrichment of two kilobases centered on the transcription start sites of 21,857 human open reading frames (NCBI RefSeq). This cost-effective approach dramatically increases the sequencing depth and resolution for nucleosome distribution measurements for the promoters of all human open reading frames.

We were interested in directly investigating nucleosome distribution differences between the CFP and CAP samples. We organized promoter architectures across all ORFs by generating k-means clustered heatmaps (k=7). These heatmaps identify nucleosomal patterns for different types of promoter organizations (Figure 1). We identified clear clusters dominated by nucleosomes at the −2, −1, +1, and +2 positions relative to the transcription start site. Most striking, however, was the concordance between all patient samples; we observed nearly identical nucleosome distribution patterns irrespective of polyp classification (Compare Fig 1, CFP and CAP). These results suggest that nucleosome distribution is invariant between CFP and CAP. We turned to the second measure of nucleosome architecture, nucleosome sensitivity.

**Figure 1.**
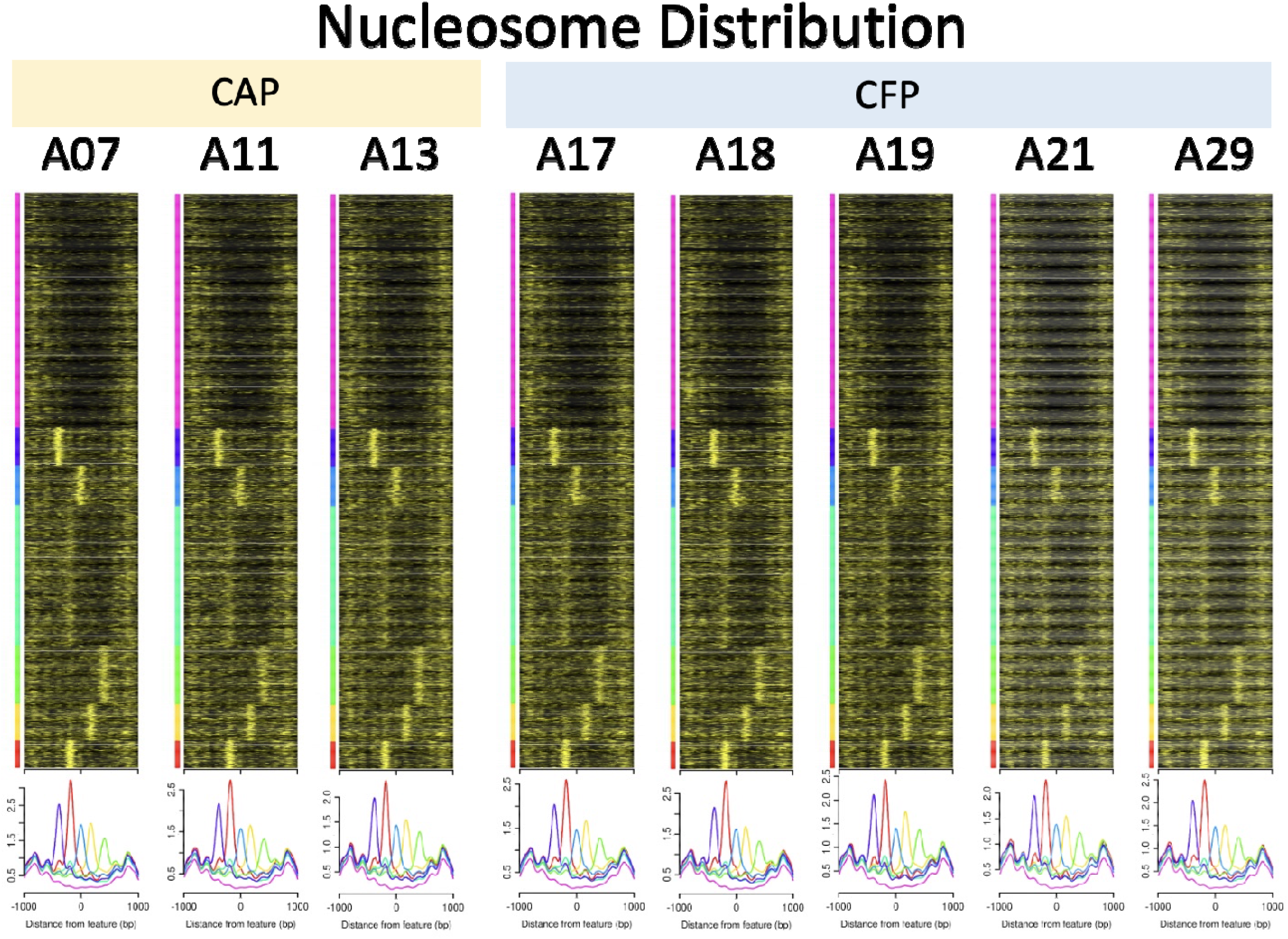
Nucleosome distribution is largely invariant between cancer free polyps and cancer adjacent polyps. Heatmaps displaying total mTSS-seq nucleosome distribution data for 2000 base pairs centered on the TSS of all human transcription start sites. Nucleosome distribution data for each patient polyp sample was organized by k-means clustering (k □ = □ 7) using the sort order of the first heatmap (CAP patient, A07). Average profiles for each cluster are shown below each individual heatmap. The left most heatmap (CAP patient A07) dictates the sort order of each map in the figure.

### Nucleosome sensitivity distinguishes cancer free and cancer associated polyps

We have recently discovered a feature of genome organization: individual nucleosomal footprints that are hypersensitive to cleavage by Micrococcal Nuclease (MNase). Using different concentrations of MNase during chromatin digestion on the resulting nucleosome position maps revealed the presence of nucleosomes susceptible to over digestion by MNase. Ours was the first laboratory to publish sensitivity maps of nucleosomes at a genome scale in multicellular organisms (12). Under light-digestion conditions, a population of genomic regions is protected by nucleosomes; however, these footprints are lost in more heavily digested states. Thus, a population of nucleosomes appears to display a biochemical property rendering them susceptible to over digestion by MNase. We refer to the regions sensitive to MNase digestion as those containing MNase Sensitive Nucleosomes (MSNs), and the less susceptible regions as MNase Resistant Nucleosomes (MRNs).

We next wanted to understand if differences between CFP and CAP could be captured by directly measuring nucleosome sensitivity. To accomplish this, we digested our patient-derived polyps with two concentrations of MNase (light and heavy, see Methods). We produced high-resolution heat maps by calculating the log2 ratio of the normalized reads mapped at 2 kb around TSSs in light over heavy MNase digests.

Clear differences in MNase sensitivity are observed within and between CFP and CAP samples (Fig. 2). It is notable that the sensitivity pattern is not maintained across samples. These results suggest that measures of nucleosome sensitivity may give insight into differences between CFP and CAP.

**Figure 2.**
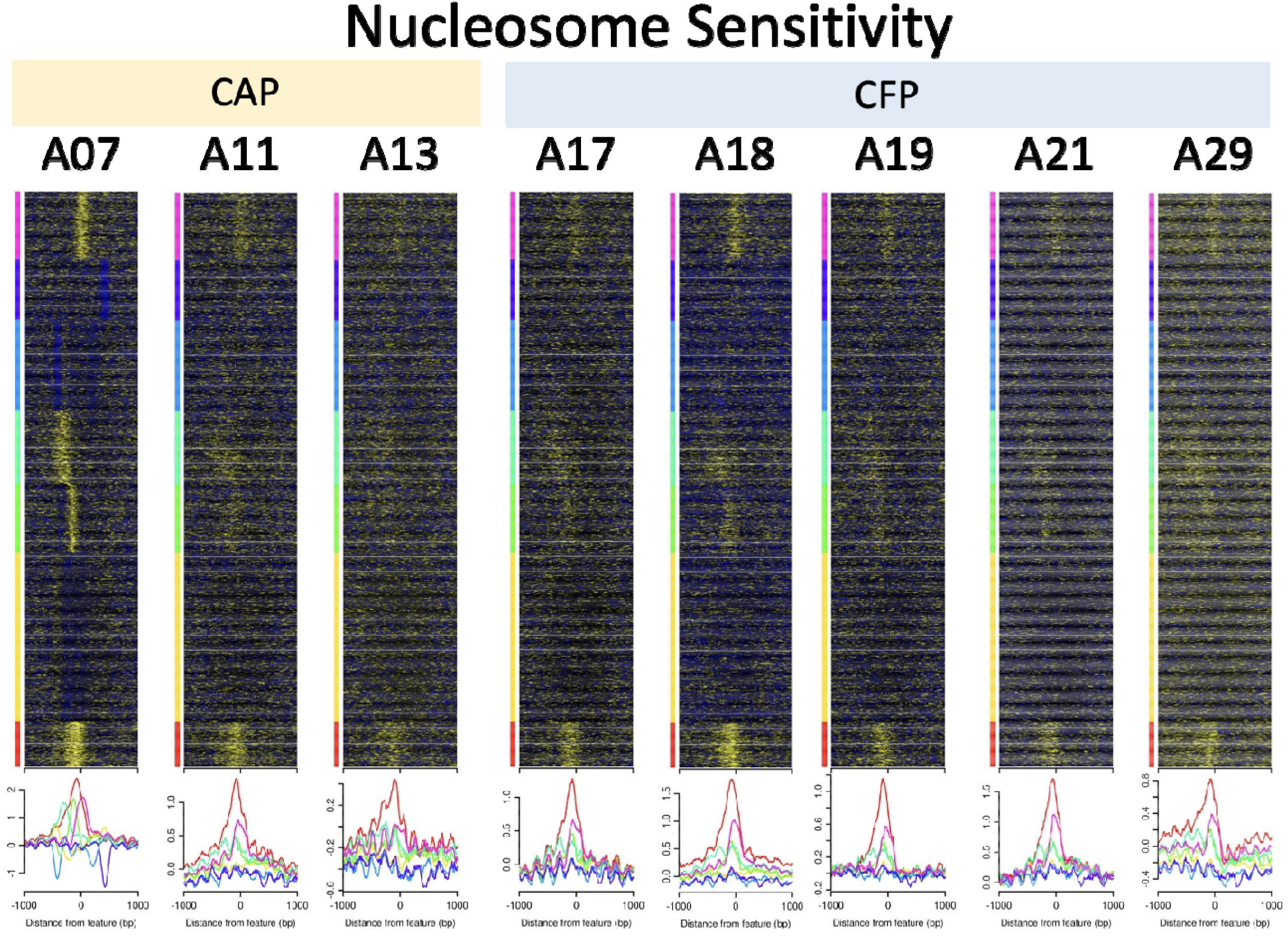
Nucleosome sensitivity is not maintained in cancer-free and cancer-associated polyps. Heatmaps displaying total mTSS-seq nucleosome sensitivity data for 2000 base pairs centered on the TSS of all human transcription start sites. Nucleosome sensitivity data for each patient polyp sample was calculated by computing the log2 ratio of reads from the light MNase digest over the reads of the heavy MNase digest. The data was organized by k-means clustering (k □ = □7) using the sort order of the first heatmap (CAP patient, A07). Average profiles for each cluster are shown below each individual heatmap. The leftmost heatmap (CAP patient A07) dictates the sort order of each map in the figure.

Nucleosome sensitivity at the +1 and -1 nucleosome positions can discriminate between CFP and CAP

The prevailing view of promoter organization suggests that the strongly positioned +1 and −1 nucleosomes, most proximal to the TSS, establish the statistical positioning of nucleosomes at the promoter, and play important roles in gene regulation (13,14). We next wanted to test the idea that the sensitivity of the -1 and +1 nucleosomes would give insights into differential promoter regulation between CFP and CAP. To accomplish this, we filtered our data at the -1 nucleosome position (−50 to -250 bp relative to TSS) or the +1 nucleosome position (+50 to +250 bp relative to TSS). We identified all -1 and +1 locations in which all patient-derived samples had a log 2 ratio of normalaized reads in light over heavy digests greater than or equal to 0.3 to be identified as sensitive, or less than -0.3 to be identified as resistant. There were two possibilities for each location CFP MRN and CAP MSN, or CFP MSN and CAP MRN. At the -1 nucleosome position, we identified 538 CFP MRN and CAP MSN and 644 CFP MSN and CAP MRN. At the +1 nucleosome position, we identified 611 CFP MRN and CAP MSN and 687 CFP MSN and CAP MRN (Fig. 3). These results suggest that MNase sensitivity has the potential to be an important analytical tool to identify functionally different promoter architectures that can discriminate CFP and CAP.

**Figure 3.**
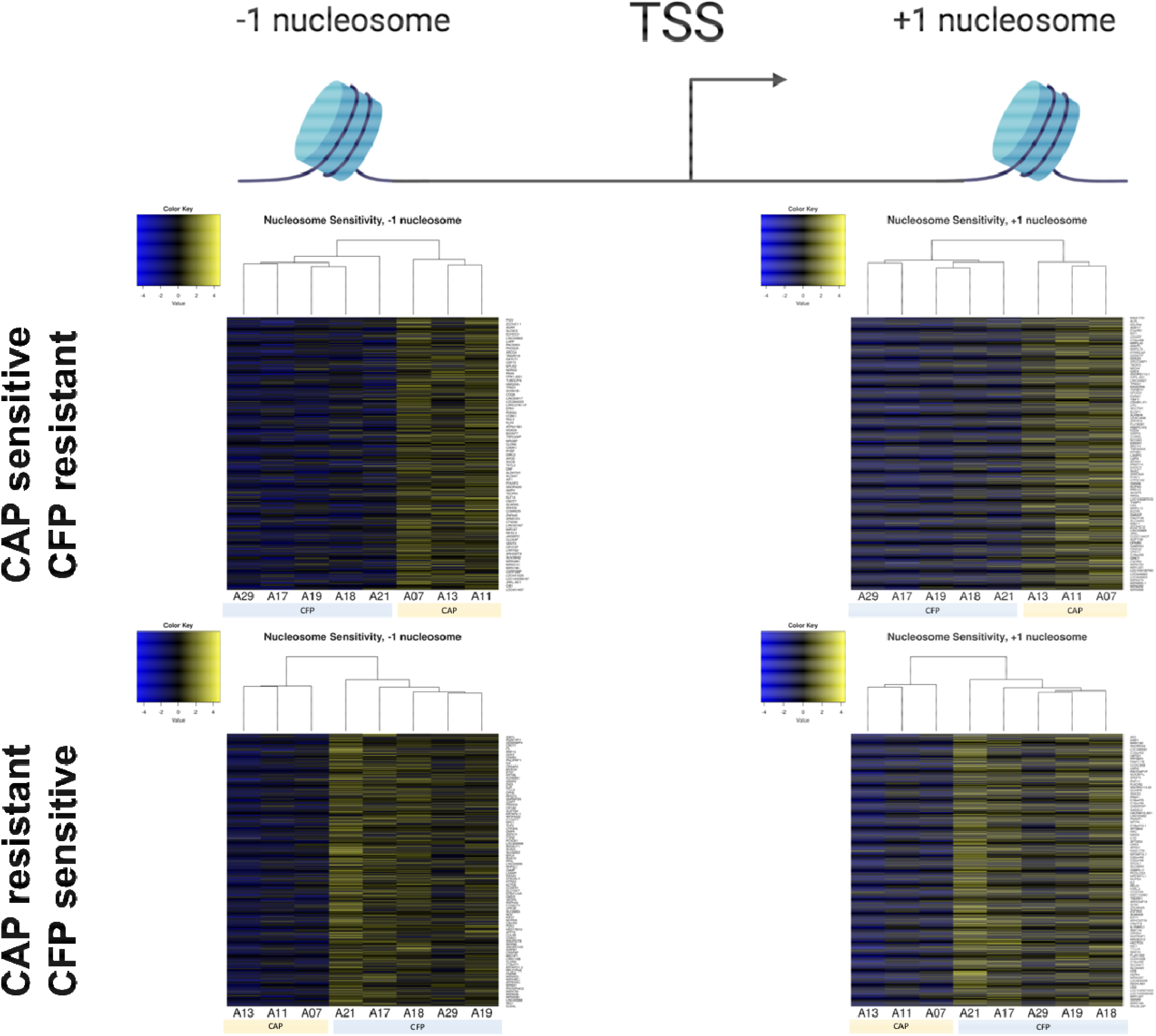
Nucleosome sensitivity at the +1 and -1 nucleosome positions can discriminate between CFP and CAP. Heatmaps showing genes that their nucleosome sensitivity at +1 nucleosome (50-250 bp downstream) and -1 nucleosome (50-250 bp upstream) shows a distinctive pattern for CAP and CFP. Columns represent patient samples, rows are genes and cells are the sensitivity value of the gene in the shown windows. Genes have been selected by comparing the sensitivity value for the same locus within and between samples in CAP and CFP polyps. The sensitivity value within sensitive group is minimum 0.3 and maximum -0.3 within resistant group. Yellow color represents sensitive nucleosome and blue color indicates resistant nucleosome.

We next wanted to further validate our MSNs and MRNs using an orthogonal method that allowed for statistical rigor. We called nucleosome-sized peaks suggestive of MSNs and MRNs using the MACS2 peak calling tool. We then calculated fold change in peak size and applied a p-value based on fragment coverage using the DESeq2 pipeline. We identified differentially sensitive regions with positive (MSN) or negative (MRN) log2 fold change and a p-value less than 0.05 (Fig. 4). Using this approach, we found 4516 MSNs and 3420 MRNs in CFPs, and 3317 MSNs and 2347 MRNs in CAPs. From this dataset, we identified four peaks that overlap between CAP MSN and CFP MRN, and three peaks that overlap between CFP MSN and CAP MRN. These results validate our initial observation that nucleosome sensitivity may be used as a tool to discriminate between CFP and CAP, and could likely serve as a powerful tool to understand the transformation process in these tissues.

**Figure 4.**
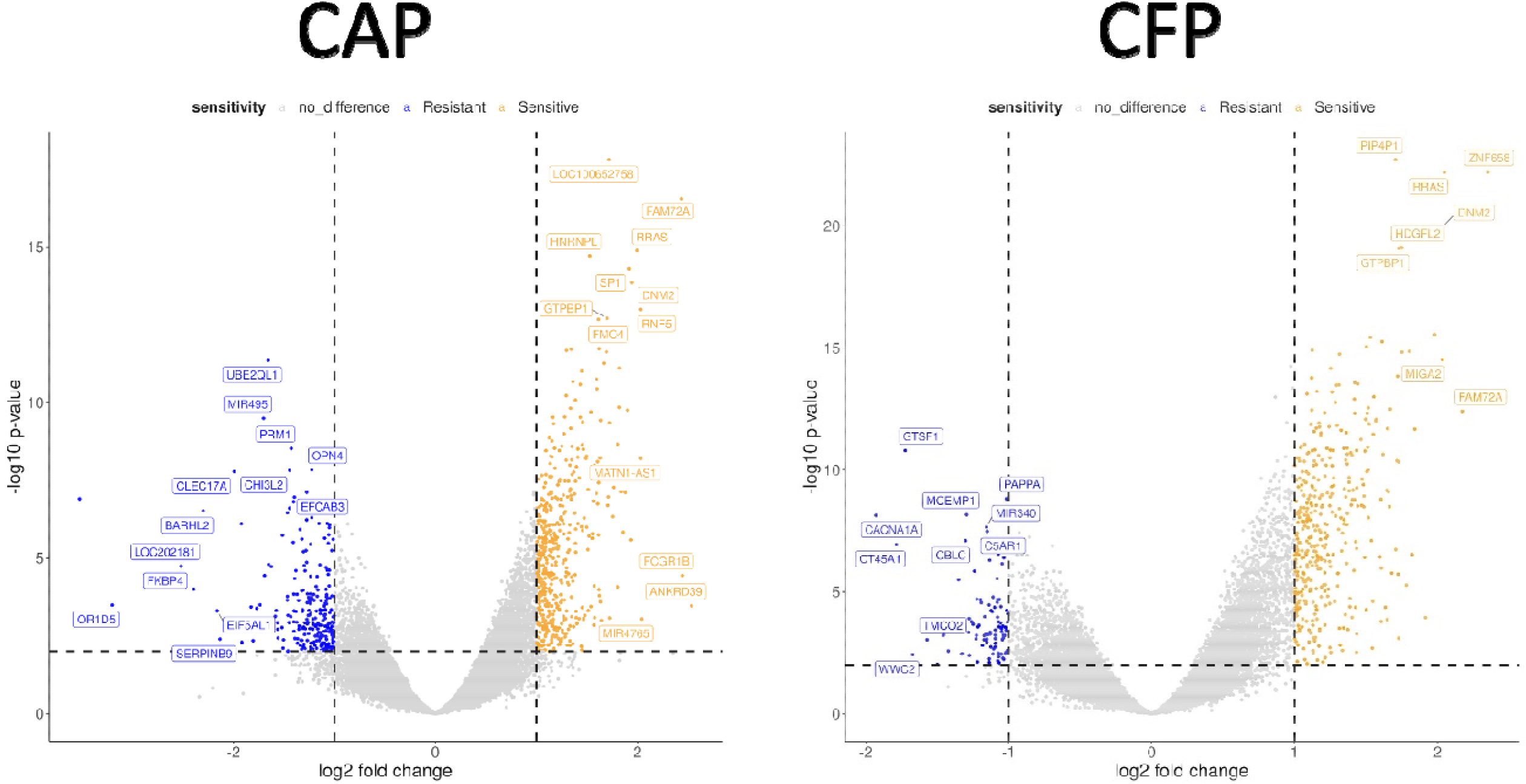
Peak analysis identifies specific genomic locations with altered nucleosome sensitivity. The volcano plots show statistically significant sensitive and resistant peaks called by MACS2. Peaks exist at 1.2 kb around the TSS of the corresponding gene and their difference is compared in light digests and heavy digests of CAP and CFP separately using DESeq2 pipeline.

### MNase sensitivity analysis identifies genes associated with transformation

Because nucleosome sensitivity could be used to identify loci and genes with differential chromatin structure between CFP and CAP, we hypothesized that these loci with different promoter architectures would give insight into specific gene functions or pathways associated with oncogenic transformation. Our analysis has identified regions with differential nucleosome sensitivity for CAP and CFP around the TSS of genes that promote malignant phenotype (Table 1). Glutaredoxin 3 (GLRX3), also known as TXNL2, Grx3, and PICOT, is involved in the maintenance of the cellular redox stated (15) and is overexpressed in pancreatic ductal adenocarcinoma (PDAC) (16), oral squamous cell carcinoma (OSCC) (17), and colon and lung carcinomas (15). GLRX3 knockdown reduces *in vitro* proliferation and *in vivo* tumor formation and could also serve as a serum biomarker for PDAC detection (16). In OSCC, GLRX3 overexpression is correlated with metastasis and the patient’s poor survival (17). Furthermore, it is shown that GLRX3 positively regulates stress induced DNA damage response by prompting ATR-dependent signaling pathways (18). Protein phosphatase 2A (PP2A) is a serine/threonine phosphatase and is involved in numerous cellular pathways, including cell proliferation, DNA damage response, and apoptosis (19,20). PP2A subunit PR130 regulates G1/S phase transition through CHK1 dephosphorylation(21). Also it is shown that PP2A is regulating the Wnt signaling (22) and regulates cell adhesion and migration by interacting with lipoma-preferred partner (23).

**Table 1.**
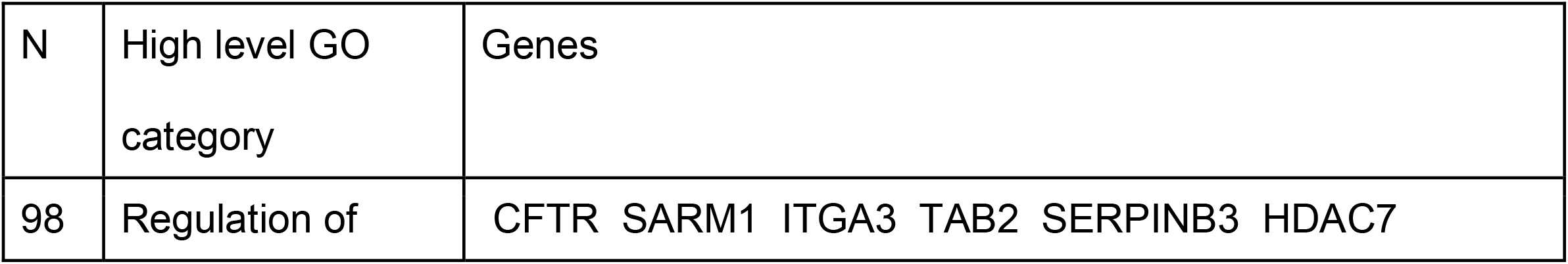
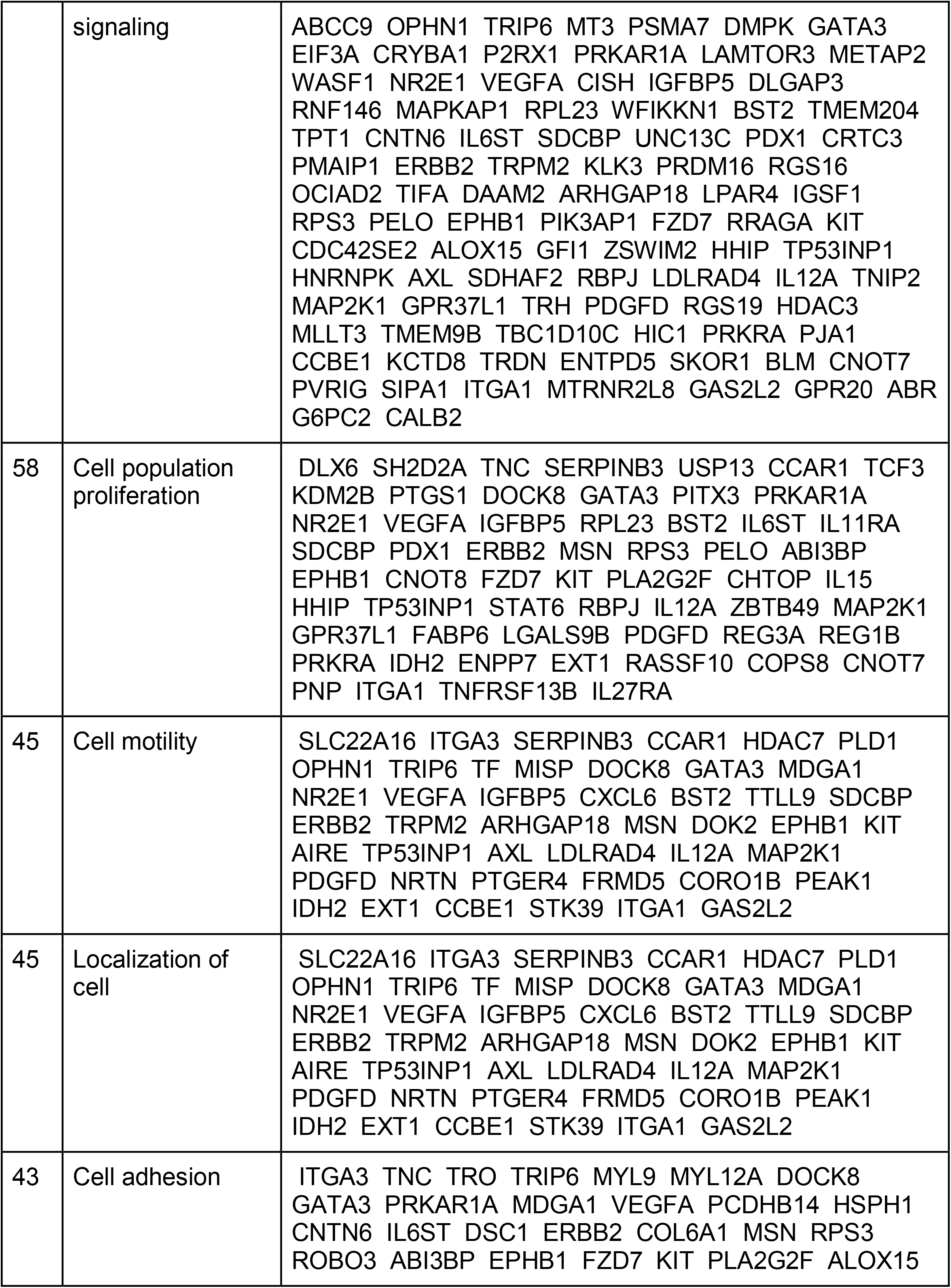
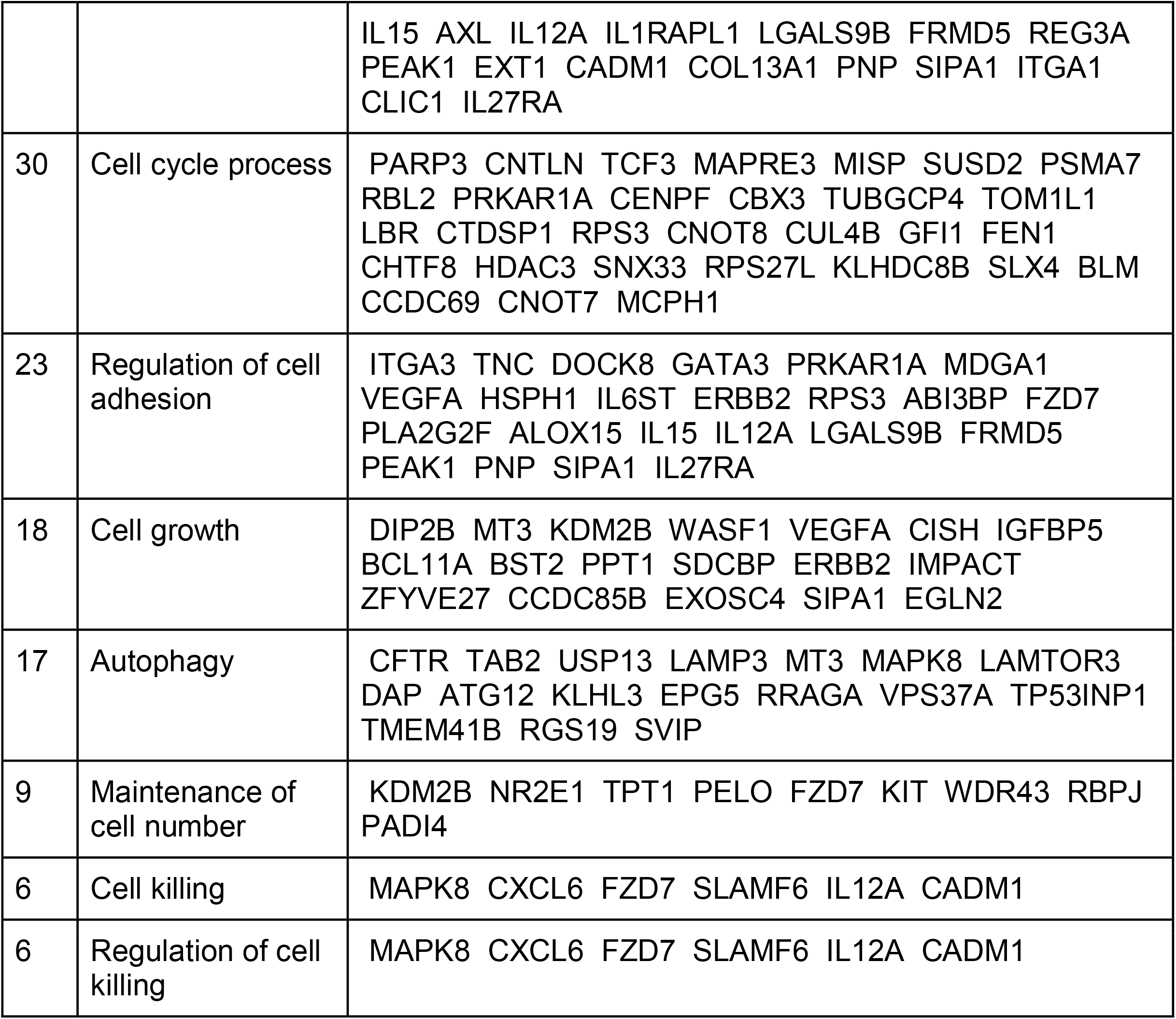
Genes with differential nucleosome sensitivity between CAP and CFP.

## DISCUSSION

The International Agency for Research on Cancer reported 19.3 million cancer cases and 10 million cancer deaths in 2020. Cancer ranks as the first or second cause of death before the age of 70 in the majority of 195 countries on earth. In addition to the sickness and death, costs reflecting treatment paid by patients, insurers, and patient time costs demonstrate the enormous economic burdens of cancer. The key to eliminating the personal and financial costs of cancer lies in the early detection of the disease. Early diagnosis relies on detecting the potentially cancerous tissue as early in disease progression as possible. Early detection is a critical public health strategy because it improves disease outcomes, increases survival, reduces complications associated with therapies, and reduces the economic burden. Early diagnosis presents a substantial challenge. There are few known early markers of cancer progression and no known markers of cancer potential. For example, colorectal cancer generally results from the oncogenic transformation of precancerous polyps in the colon or rectum.Polyps are endoscopically or surgically removed, and the patient is followed with colonoscopy at regular intervals for 3-5 years. There are currently no clinical tests or screening to identify biomarkers that classify a polyp as likely to transform. Our results suggest that the identification of MSN and MRN reveals an important set of biomarkers that could be used to classify the transformation potential of resected polyps found during routine colonoscopy screens. We suggest that the resected polyps could be used in nucleosome sensitivity assays to provide patients and physicians with the earliest possible information on the transformation potential of a polyp.

## MATERIALS AND METHODS

### Patient sample characteristics and tissue preparation

All tissues were collected at Mayo Clinic between 2000-2016 through an IRB approved Biobank for Gastrointestinal Health Research [BGHR] (IRB 622-00), as described previously in Druliner et al (8). Polyp tissues with adjacent tumor and normal colonic epithelium full thickness specimens at least 8 cm from the polyp/tumor margin were harvested following surgical resection and snap frozen in liquid nitrogen and maintained in a -80 freezer. Cancer free polyps and normal colonic epithelium at least 8 cm from the polyp were collected at the time of colonoscopic resection. We utilized cancer adjacent polyp (CAP) and cancer free polyp (CFP) cases. Cancer adjacent polyps (CAPs) were matched to the cancer free polyps (CFPs) based on polyp size (categorical size: 1 to 2 cm, 2-5 cm and > 5 cm); histology (villous features) and degree of dysplasia. All polyps presented in this study are adenomatous polyps with villous features (tubulovillous or villous) and with low grade dysplasia only. All CAP and CFP cases exclude subjects with a prior history of any malignancy; a family history of Lynch syndrome or Familial Adenomatous Polyposis (FAP); any other syndrome associated with hereditary CRC or inflammatory bowel disease. All tissue used in this study was removed prior to neoadjuvant/adjuvant therapy. CAPs were cases in which histological review of the surgically resected CRC showed the residual polyp of origin (RPO) in direct contiguity to the cancer. Tissues were macro-dissected using a hematoxylin and eosin (H&E) guide that was used to mark areas of normal epithelium, polyp, or cancer by a pathologist.

### MNase cleavage and purification of mononucleosomal and subnucleosomal DNA

Five 10um cryostat slices of patient polyps were digested for 5 min at 37°C with a titration of MNase: 20 units/mL and 200 unit/mL of MNase (Worthington Biochemical Corp.) in MNase cleavage buffer (4 mM MgCl2, 5 mM KCl, 50 mM Tris-Cl (pH 7.4), 1 mM CaCl2, 12.5% glycerol). The MNase digestion reactions were stopped with 50 mM EDTA. Next, the protein-DNA crosslinks were reversed by treating the MNase-digested nuclei with 0.2 mg/mL proteinase K and 1% sodium dodecyl sulfate, and incubating overnight at 60°C.

The samples were then on a 2% agarose gel. Following the separation of the DNA fragments, mononucleosomally-sized and subnucleosomal-sized fragments (<200 bp) were isolated from the agarose gel, and the DNA was purified by freeze and squeeze.

#### Mononucleosomal and subnucleosomal DNA library preparation

The NEBNext® Ultra™ DNA Library Prep Kit for Illumina® (NEB #E7370S/L) was used to prepare DNA sequencing libraries. Briefly, 30 ng of DNA was end-prepped and adaptors were ligated. The adaptor-ligated DNA was cleaned up using AMPure XP beads to remove any unwanted ligated products. The adapted sequences were indexed using NEB Unique Dual indices by eight cycles of PCR. Unincorporated indices were removed the indexed DNA using AMPure XP beads. The libraries were quality-checked using Agilent High-Sensitivity Tape Station. The samples had a mean size of 383 basepairs.

### Solution-based sequence-capture of indexed libraries

Utilizing our custom-designed Roche Nimblegen SeqCap EZ Library SR, we sequence-captured the indexed fragments within the two-kb window surrounding all human TSSs. Following the sequence-capture, we PCR amplified our sequence-captured fragments using TruSeq primers 1 and 2 (AATGATACGGCG ACCACCGAGA and CAAGCAGAAGACGGCATACGAG, respectively).

### Illumina paired-end sequencing and analysis

The samples were loaded at 12 pM onto a NovaSeq6000 for paired end 150 sequencing. The reads were demultiplexed using the Casava Software, and the library adaptors were removed using the trimmomatic software.

### Alignment and data processing bioinformatics

Casava software was used to demultiplex the indices in each lane. Illumina adapters were clipped from reads with trimmomatic and aligned to the hg19 human genome assembly with bowtie2 2.1.0 with default parameters. Unpaired and non-uniquely-mapped reads were discarded with samtools. Individual nucleosome footprints were extracted from BAM files with bedtools 2.17. Nucleosome occupancy profiles were obtained by calculating the fragments per million that mapped at each base-pair in the probed regions with bedtools. Nucleosome dyad frequencies (midpoints) were obtained by calculating the sum of nucleosome dyads (fragment centers) in 60-bp windows at a 10-bp step-size with bedtools. Data were subsequently processed in R 2.15.1. Data was uploaded to the UCSC genome browser for further analysis (http://genome.ucsc.edu).

### Nucleosome distribution maps were prepared using BAM files and of bedtools 2.17

100 bp from the center of the mapped reads was computationally selected for the fragments that were larger than 100 bp. Nucleosome distributions were normalized to fragments per million in the 2kb surrounding each TSS. Further analysis of the nucleosome distributions R utilized our lab-developed software, GENMAT run in the R environment, R 2.15.1.

